# Two variance component model improves genetic prediction in family data sets

**DOI:** 10.1101/016618

**Authors:** George Tucker, Po-Ru Loh, Iona M MacLeod, Ben J Hayes, Michael E Goddard, Bonnie Berger, Alkes L Price

## Abstract

Genetic prediction based on either identity by state (IBS) sharing or pedigree information has been investigated extensively using Best Linear Unbiased Prediction (BLUP) methods. Such methods were pioneered in the plant and animal breeding literature and have since been applied to predict human traits with the aim of eventual clinical utility. However, methods to combine IBS sharing and pedigree information for genetic prediction in humans have not been explored. We introduce a two variance component model for genetic prediction: one component for IBS sharing and one for approximate pedigree structure, both estimated using genetic markers. In simulations using real genotypes from CARe and FHS family cohorts, we demonstrate that the two variance component model achieves gains in prediction *r*^2^ over standard BLUP at current sample sizes, and we project based on simulations that these gains will continue to hold at larger sample sizes. Accordingly, in analyses of four quantitative phenotypes from CARe and two quantitative phenotypes from FHS, the two variance component model significantly improves prediction *r*^2^ in each case, with up to a 20% relative improvement. We also find that standard mixed model association tests can produce inflated test statistics in data sets with related individuals, whereas the two variance component model corrects for inflation.

**Author Summary:** Genetic prediction has been well-studied in plant and animal breeding and has generated considerable recent interest in human genetics, both in family data sets and in population cohorts. Many prediction studies are based on the widely used Best Linear Unbiased Prediction (BLUP) approach, which performs a mixed model analysis using a genetic relationship matrix that is either estimated from genotype data—thus measuring identity-by-state (IBS) sharing—or obtained from family pedigree information. We show here that a substantial improvement in prediction accuracy in family data sets can be obtained by jointly modeling both IBS sharing and approximate pedigree structure, both estimated using genetic markers, using separate variance components within a two variance component mixed model. We demonstrate the performance of this model in simulations and real data sets. We also show that previous mixed model association methods suffer from inflated test statistics in family data sets due to their failure to account for the different heritability parameters corresponding to IBS sharing vs. pedigree relatedness. Our two variance component model provides a solution to this problem without compromising statistical power.

## Introduction

Mixed linear models (MLM) are widely used for genetic prediction and association testing in genome-wide association studies (GWAS). In prediction, MLM produce best linear unbiased predictions; BLUP and its extensions were first developed in agricultural genetics [1–4] and have since been applied to human genetics [5–10]. In association testing, MLM model relatedness and population stratification, correcting for confounding and increasing power over linear regression (essentially by testing association of the residual from BLUP prediction) [11–16]. Mixed model methods harness information from either genetic markers (IBS sharing) or known pedigree relationships. Recent work on estimating components of heritability [17] has demonstrated the advantages of a model with two variance components: one component for IBS sharing (corresponding to SNP-heritability 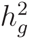 [18, 19]) and one for approximate pedigree structure, estimated via IBS sharing above a threshold (corresponding to total narrow-sense heritability *h*^2^ [20]). However, the potential advantages of this model for genetic prediction (or mixed model association) have not been explored.

Through systematic simulations and analyses of quantitative phenotypes in the Candidate-gene Association Resource (CARe) [21] and Framingham Heart Study (FHS) [22, 23] cohorts, we show that the two variance component model improves prediction *r*^2^ over single variance component (standard BLUP) methods. Our simulations demonstrate that this improvement is achieved both at current sample sizes and for larger samples, and our analyses of real CARe and FHS phenotypes confirm relative improvements in prediction *r*^2^ of up to 20%. We also consider the situation in which phenotypes are available for ungenotyped individuals that are related to the genotyped cohort (e.g., family history [24, 25]) and show that leveraging this additional information for genetic prediction within a two variance component model achieves similar gains.

Additionally, we investigate the utility of the two variance component model for association testing. We evaluate the standard prospective MLM association statistic [15] in the context of familial relatedness and observe inflation of test statistics over a range of simulation parameters, contrary to previous findings [11, 13–15, 26]. We show that the two variance component model substantially reduces the inflation in simulations and in GWAS of CARe and FHS phenotypes.

## Results

### Overview of Methods

We use the two variance component model described in previous work on estimating components of heritability [17]. The first variance component is the usual genetic relationship matrix (GRM) computed from genetic markers (corresponding to 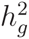) [18]. The second variance component is a thresholded version of the GRM in which pairwise relationship estimates smaller than a threshold *t* are set to zero, the idea being to capture strong relatedness structure similar to a pedigree relationship matrix. Explicitly modeling relatedness in this way allows the two variance component mixed model to capture additional heritability from untyped SNPs (corresponding to 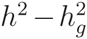) [17]. We used the two variance component model to compute genetic predictions via BLUP and test associations using a Wald statistic [1, 11, 27]. (We note that best linear unbiased prediction, BLUP, is a general method for prediction that can be applied once a covariance model has been established, whether from one or many variance components. We will therefore use “standard BLUP” to refer to BLUP using the GRM as a single variance component, and we will use “BLUP” to more generally refer to BLUP with any number of variance components.) We further developed methods to treat the case in which phenotypes for ungenotyped relatives are available; briefly, our approach uses pedigree information to impute the missing information [28]. Full details are provided in Materials and Methods and Text S1.

### Genetic prediction: simulations

To analyze the predictive power of the two variance component model, we simulated phenotypes based on genotypes from the CARe and FHS data sets as described in Materials and Methods. In each simulation, we used the following procedure to measure prediction accuracy of BLUP using the standard GRM as a single variance component, BLUP using the thresholded GRM as a single variance component, and BLUP using the two variance component model. First, we randomly split the data set, setting aside 90% of the individuals for training and 10% for testing. We then used each BLUP method to predict held-out test phenotypes using the training samples to estimate genetic effects, and we calculated *r*^2^ between the predicted phenotypes and the true genetic components of the simulated phenotypes (i.e., eliminating the added noise) on the test samples. (We chose to compute *r*^2^ as it is a very widely used metric for assessing prediction accuracy [2–4, 6, 7, 9]; however, other metrics such as mean square error are also sometimes used [5].) We call this quantity “prediction *r*^2^(*g*)”; on average, prediction *r*^2^(*g*) is 1*/h*^2^ times as large as standard prediction *r*^2^, i.e., *r*^2^ computed to simulated phenotypes that include both genetic and noise components. Relative performance of prediction methods is the same (on average) whether measured with prediction *r*^2^ or prediction *r*^2^(*g*).

The two variance component model provided significant increases in *r*^2^(*g*) over both standard BLUP and BLUP using the thresholded GRM alone (Table 1), and the improvements were consistent across simulation replicates (Fig. S1). We observed much larger prediction *r*^2^(*g*) values (across all methods) for the FHS simulations than the CARe simulations, as expected given the much greater number of close relatives in the FHS data set (18,415 pairs of individuals with genetic relatedness *>*0.2 among 7,476 FHS individuals vs. 4,954 pairs among 8,367 CARe individuals). However, the relative improvements achieved by the two variance component model were fairly similar in these two distinct pedigree structures, and importantly, increasing values of *N/M* (mimicking larger sample sizes) also yielded similar relative improvements (Table 1). We also observed that the heritability parameter estimated by the standard mixed model was intermediate to 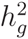 and *h*^2^, whereas the two variance component model accurately estimated 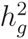 and 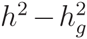 (Table S1), as expected in samples with related individuals [17].

**Table 1.** Prediction accuracy for simulations using CARe and FHS genotypes. Phenotypes were simulated to have *h*^2^ = 0.5, 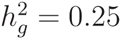, and prediction *r*^2^(*g*) was measured using a random 90% of samples as training data and the remaining 10% as test data. Reported values are mean prediction *r*^2^(*g*) and s.e.m. over 100 independent simulations (in which phenotypes were re-simulated and train/test splits resampled). BLUP w/ thresh. denotes BLUP prediction using the thresholded relationship matrix instead of the standard approach of using the GRM (denoted simply “BLUP”). Prediction *r*^2^(*g*) denotes *r*^2^ between predicted phenotypes and true genetic components of the simulated phenotypes.

| Observed SNPs | Prediction $r^2(g)$ | | |
| --- | --- | --- | --- |
|  | BLUP | BLUP w/ thresh. | 2 VC BLUP |
| Chrom. 2 - 22 | 0.062 (0.002) | 0.061 (0.002) | 0.071 (0.002) |
| Chrom. 3 - 6 | 0.084 (0.002) | 0.063 (0.002) | 0.094 (0.002) |
| Chrom. 3 - 4 | 0.098 (0.002) | 0.059 (0.002) | 0.108 (0.002) |

(b) FHS genotypes
| Observed SNPs | Prediction $r^2(g)$ | | |
| --- | --- | --- | --- |
|  | BLUP | BLUP w/ thresh. | 2 VC BLUP |
| Chrom. 2 - 22 | 0.225 (0.003) | 0.225 (0.003) | 0.238 (0.003) |
| Chrom. 3 - 6 | 0.246 (0.003) | 0.230 (0.003) | 0.269 (0.003) |
| Chrom. 3 - 4 | 0.263 (0.003) | 0.231 (0.003) | 0.291 (0.003) |

Finally, we assessed the potential performance of the two variance component approach at very large values of *N/M* (up to 100) by simulating both genotypes and phenotypes (Materials and Methods). (We note that human genotyping arrays typically contain ≈ 60,000 independent SNPs [15, 29], so *N/M*=8 in this simulation corresponds to a data set the size of UK Biobank, *N*=500,000; see Web Resources.) In these simulations, we continued to observe gains using the two variance components approach; two variance component prediction *r*^2^ exceeded 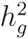 for very large *N*, whereas standard BLUP prediction *r*^2^ was limited to less than 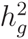 (Fig. S2).

### Genetic prediction: real phenotypes

Next, we evaluated the prediction accuracy of each method on CARe phenotypes—body mass index (BMI), height, low density lipoprotein cholesterol (LDL), and high density lipoprotein cholesterol (HDL)—and for FHS phenotypes—height and BMI. We adjusted phenotypes for age, sex, study center (for CARe phenotypes), and the top 5 principal components. (The complexities of the impact of ancestry on genetic prediction are discussed in ref. [30].) To measure performance, we created 100 independent random 90/10 splits of the data set as before and calculated *r*^2^ between predicted and true phenotypes on the test samples of each split. We observed that for all phenotypes, the two variance component model increased prediction accuracy over both single variance component BLUP approaches, with a maximum relative improvement of 20% for height (Table 2a,b); this improvement was consistent across different train/test splits (Figure S3). As in our simulations, we observed larger absolute prediction *r*^2^ in FHS than CARe due to strong relatedness (consistent with ref. [6]), and we observed that the heritability parameter estimated by the standard mixed model was intermediate to the heritability parameters 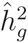 and 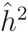 estimated by the two variance component model (Table S2).

**Table 2.** Prediction accuracy for CARe and FHS phenotypes. CARe prediction using genome-wide significant SNPs as fixed effect covariates FHS prediction using genome-wide significant SNPs as fixed effect covariates Prediction *r*^2^ values are means over 100 random 90*/*10 train/test data splits. Relative performance values reported are ratios of means minus 1; standard errors are estimatevd as standard deviations of per-split differences in *r*^2^ (over the random 10% test sets) divided by 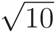 (to account for the 10x larger sample size of the full data set; see Materials and Methods). BLUP w/ thresh. denotes BLUP prediction using the thresholded relationship matrix instead of the standard approach of using the GRM (denoted simply “BLUP”).

| Phenotype | Prediction $r^2$ | | | Prediction $r^2$ relative to BLUP (s.e.) | |
| --- | --- | --- | --- | --- | --- |
|  | BLUP | BLUP w/ thresh. | 2 VC BLUP | BLUP w/ thresh. | 2 VC BLUP |
| BMI | 0.023 | 0.027 | 0.029 | +14% (9%) | +18% (5%) |
| height | 0.063 | 0.067 | 0.079 | +5% (5%) | +20% (3%) |
| LDL | 0.017 | 0.017 | 0.019 | +2% (15%) | +11% (5%) |
| HDL | 0.034 | 0.032 | 0.038 | -7% (10%) | +11% (4%) |

**(b) FHS prediction**
| Phenotype | Prediction $r^2$ | | | Prediction $r^2$ relative to BLUP (s.e.) | |
| --- | --- | --- | --- | --- | --- |
|  | BLUP | BLUP w/ thresh. | 2 VC BLUP | BLUP w/ thresh. | 2 VC BLUP |
| BMI | 0.103 | 0.104 | 0.107 | +1.0% (2.3%) | +3.5% (1.2%) |
| height | 0.344 | 0.342 | 0.354 | -0.7% (1.1%) | +2.9% (0.5%) |

**(c) CARE prediction using genome-wide significant SNPs as fixed effect covariates**
| Phenotype | Prediction $r^2$ | | | Prediction $r^2$ relative to BLUP (s.e.) | |
| --- | --- | --- | --- | --- | --- |
|  | BLUP | BLUP w/ thresh. | 2 VC BLUP | BLUP w/ thresh. | 2 VC BLUP |
| BMI | 0.023 | 0.026 | 0.028 | +14% (9%) | +19% (5%) |
| height | 0.063 | 0.066 | 0.078 | +5% (5%) | +20% (3%) |
| LDL | 0.038 | 0.039 | 0.041 | +3% (6%) | +6% (2%) |
| HDL | 0.051 | 0.049 | 0.055 | -4% (6%) | +7% (3%) |

**(d) FHS prediction using genome-wide significant SNPs as fixed effect covariates**
| Phenotype | Prediction $r^2$ | | | Prediction $r^2$ relative to BLUP (s.e.) | |
| --- | --- | --- | --- | --- | --- |
|  | BLUP | BLUP w/ thresh. | 2 VC BLUP | BLUP w/ thresh. | 2 VC BLUP |
| BMI | 0.105 | 0.107 | 0.109 | +1.2% (2.3%) | +3.5% (1.2%) |
| height | 0.344 | 0.341 | 0.354 | -0.8% (1.1%) | +2.8% (0.5%) |

For phenotypes with a small number of large effect loci, methods that explicitly model a non-infinitesimal genetic architecture can have substantially better prediction accuracy than standard BLUP [2]. A two variance component approach could be combined with such models, and as an initial exploration of this approach, we examined a non-infinitesimal extension of two variance component BLUP in which we included large-effect loci as fixed effect covariates [8]. Explicitly, we first identified genome-wide-significant SNPs (*p <* 5 × 10^−8^) according to a two variance component mixed model association statistic. (As we show below, the standard MLM statistic is miscalibrated in scenarios with pervasive relatedness, precluding its use.) We then added these SNPs as fixed effect covariates in all of the models we previously compared and recomputed predictions (Table 2c,d). Including large-effect loci resulted in substantial improvements in prediction *r*^2^ achieved by each model for the CARe HDL and LDL phenotypes (Table 2c), both of which are known to have several large-effect loci [31]. As before, for all phenotypes, we observed an increase in *r*^2^ when using the two variance component model. We expect that the two variance component model will provide similar improvements in prediction *r*^2^ if incorporated in more sophisticated non-infinitesimal models (e.g., [3, 5]).

Additionally, we explored the scenario in which some phenotypes are available for ungenotyped relatives of genotyped individuals. We simulated data with ungenotyped individuals by randomly masking the genotypes of 25% of the training individuals. Results on simulated and real phenotypes using this masking are broadly consistent with results reported above with all individuals typed (Tables S3–S6).

### Association testing

We next compared mixed model association testing using the two variance component approach to standard MLM association testing [12, 15] in data sets with related individuals, measuring calibration and power for each method. We began by running a suite of tests using simulated genotypes and phenotypes, systematically varying the number of related individuals, the degree of relatedness, the number of markers in the genome, and the heritability of the simulated trait (see Materials and Methods). Each simulation included both causal SNPs and “null SNPs,” i.e., SNPs with no phenotypic effect. For null SNPs, Wald statistics computed by mixed model association tests follow a 1 d.o.f. chi-squared distribution assuming the mixed model accurately models the phenotypic covariance. If the mixed model does not accurately model the covariance, as we expect for phenotypes with 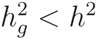 in data sets containing relatedness, then the distribution of association statistics at null SNPs is miscalibrated, i.e., approximately follows a scaled 1 d.o.f. chi-squared [32]. We therefore measured calibration of MLM association methods by computing the mean Wald statistic over null SNPs. We measured power by dividing the mean Wald statistic over causal SNPs by the mean Wald statistic over null SNPs. Computing the ratio in the latter benchmark ensured that all methods, including those susceptible to inflation of test statistics, were equally calibrated before comparing power.

Contrary to previous work suggesting that mixed models fully correct for relatedness [11, 13– 15, 26], we found that for many parameter settings, standard MLM association analysis produced significantly inflated test statistics (up to 11% inflation, increasing with trait heritability, sample size, and extent of relatedness; Figure 1). In contrast, introducing a second variance component—either the thresholded GRM (Figure 1) or the true pedigree (Fig. S4)—nearly eliminated the inflation. For all parameter settings, we observed that the two variance component model maintained or slightly increased power compared to standard MLM association (Fig. S4).

**Figure 1.**
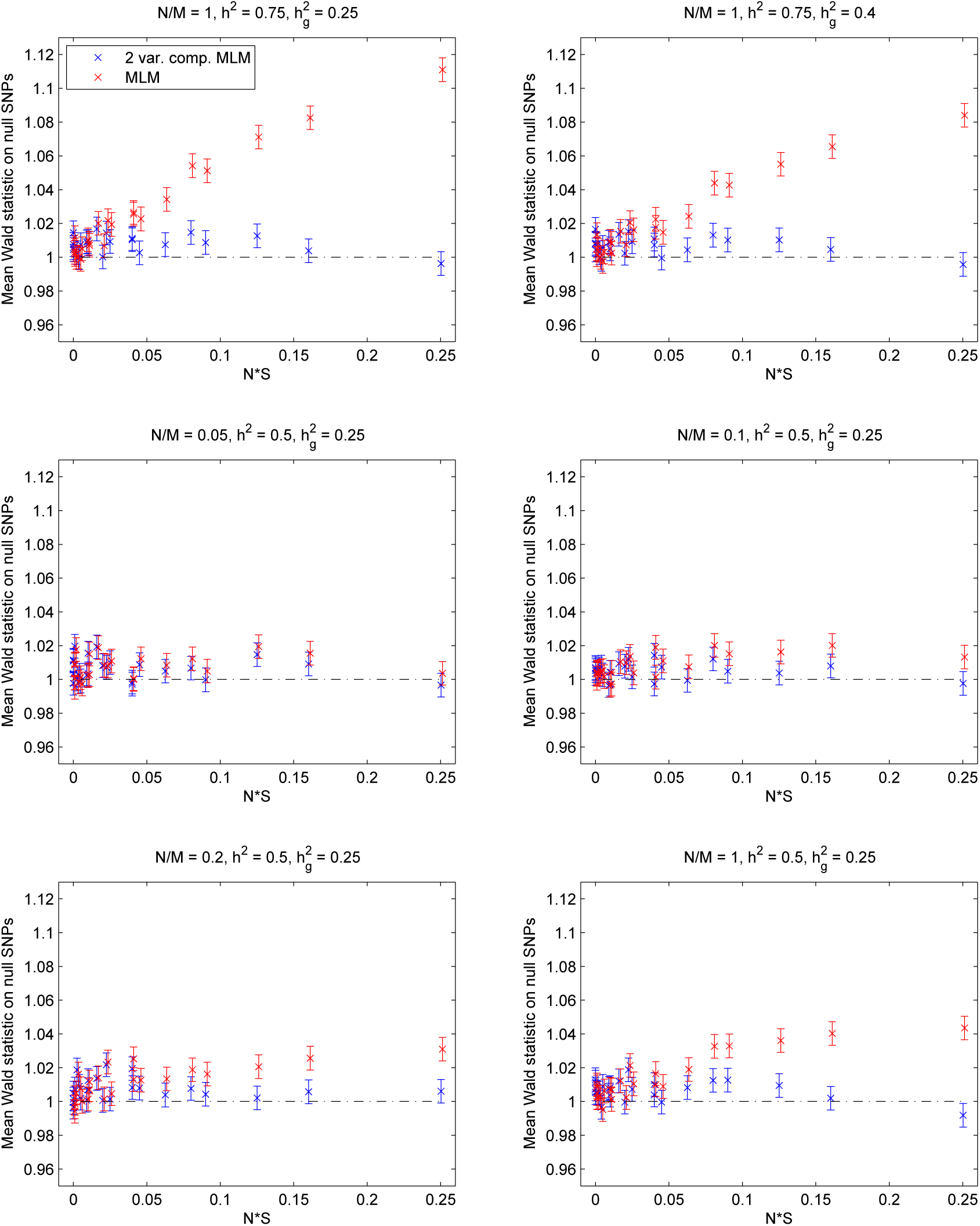
Calibration of standard and two-variance-component mixed model association statistics on simulated genotypes and phenotypes. We computed mean Wald statistics over null SNPs using the standard mixed model association test (MLM) and a two variance component model (2 var. comp. MLM) using GRM and thresholded GRM (i.e., approximate pedigree) components. Each panel shows results from a set of simulations with selected values of the simulation parameters *N/M*, *h*^2^, and 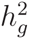. The set of simulations contained within each panel varies one additional parameter, *NS*, which measures the amount of relatedness in the simulated data. (*S* denotes the average squared off-diagonal entry of the pedigree relationship matrix.) Plotted values are mean Wald statistics and s.e.m. over 100 simulations.

Next, we simulated phenotypes based on genotypes from the CARe and FHS data sets (Materials and Methods). Consistent with the previous simulations, standard MLM association produced inflated statistics whereas the two variance component model alleviated inflation (Tables 3 and S7). Importantly, these results suggest that the levels of relatedness that are required for inflation are present in real data sets.

**Table 3.** Calibration of standard and two-variance-component mixed model association statistics in CARe and FHS simulations. Mean Wald statistics on candidate null SNPs for simulations with CARe or FHS genotypes and a trait with *h*^2^ = 0.5, 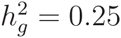. Reported values are means and s.e.m. over 100 simulations. The two variance component model selected the specified threshold (*t*) to estimate the relatedness matrix. In simulations using only SNPs on chromosome 22 to compute GRMs, we observed slight inflation using the two variance component model; given the large thresholds (*t >* 0.25) chosen by the model in these scenarios, we hypothesize that the number of SNPs was too small to distinguish relatedness from noise in the GRM, causing an incomplete correction. For corresponding Type I error at different *α* levels, see Table S7.

| Observed SNPs | # of SNPs<br>($M$) | Standard mixed model<br>Mean Wald | Two variance components | |
| --- | --- | --- | --- | --- |
| | | | Mean Wald | Threshold ($t$) |
| Chrom. 3 - 22 | 615,445 | 1.013 (0.002) | 1.000 (0.002) | 0.024 |
| Chrom. 3 - 6 | 195,333 | 1.024 (0.002) | 1.002 (0.002) | 0.051 |
| Chrom. 3 - 4 | 99,690 | 1.028 (0.002) | 1.003 (0.002) | 0.081 |
| Chrom. 22 | 9,713 | 1.036 (0.002) | 1.014 (0.002) | 0.387 |

(b) FHS genotypes
| Observed SNPs | # of SNPs<br>($M$) | Standard mixed model<br>Mean Wald | Two variance components | |
| --- | --- | --- | --- | --- |
| | | | Mean Wald | Threshold ($t$) |
| Chrom. 3 - 22 | 346,005 | 1.032 (0.003) | 1.003 (0.003) | 0.021 |
| Chrom. 3 - 6 | 110,203 | 1.071 (0.003) | 1.008 (0.003) | 0.040 |
| Chrom. 3 - 4 | 55,480 | 1.097 (0.003) | 1.014 (0.003) | 0.055 |
| Chrom. 22 | 5,277 | 1.189 (0.004) | 1.055 (0.003) | 0.258 |

Finally, we analyzed MLM association statistics for the CARe and FHS phenotypes (adjusted for covariates as before). Because we do not know the identity of causal and null SNPs in this case, we calculated the average Wald statistic over all SNPs using leave-one-chromosome-out analysis [15, 33], noting that we expect the statistics to be slightly larger than 1 due to polygenicity [15,29]. Consistent with simulations, the average Wald statistics were higher for standard MLM association than the two variance component method, suggesting that standard MLM statistics are slightly inflated, with an up to 1.05-fold inflation in FHS data (Table 4).

**Table 4.** Calibration of standard and two-variance-component mixed model association statistics for CARe and FHS phenotypes. We report the number of individuals *N* phenotyped for each trait and the mean Wald statistics and heritability parameters computed by the standard and two-variance-component mixed models (averaged over 22 leave-one-chromosome-out runs).

| Phenotype | $N$ | Standard mixed model | | Two variance components | | |
| --- | --- | --- | --- | --- | --- | --- |
| | | Mean Wald | $\hat{h}_g^2$ | Mean Wald | $\hat{h}_g^2$ | $\hat{h}^2$ |
| BMI | 7987 | 1.044 | 0.35 | 1.029 | 0.17 | 0.46 |
| height | 7988 | 1.110 | 0.73 | 1.080 | 0.38 | 0.95 |
| LDL | 4965 | 1.030 | 0.32 | 1.021 | 0.18 | 0.44 |
| HDL | 5184 | 1.054 | 0.50 | 1.037 | 0.26 | 0.66 |

(b) FHS phenotypes
| Phenotype | $N$ | Standard mixed model | | Two variance components | | |
| --- | --- | --- | --- | --- | --- | --- |
| | | Mean Wald | $\hat{h}_g^2$ | Mean Wald | $\hat{h}_g^2$ | $\hat{h}^2$ |
| BMI | 7476 | 1.060 | 0.43 | 1.032 | 0.21 | 0.47 |
| height | 7476 | 1.126 | 0.81 | 1.070 | 0.39 | 0.87 |

## Discussion

We have shown that a mixed model with two variance components, one modeling genetic effects of typed SNPs and the other modeling phenotypic covariance from close relatives, offers increased prediction accuracy over standard BLUP and corrects miscalibration of standard mixed model association analysis in human data sets containing strong relatedness. For current sample sizes and levels of relatedness, the absolute increase in prediction accuracy is modest (similar to other recent work on improving prediction accuracy for human complex traits [5, 7–10], in contrast to agricultural traits [2–4]) and the inflation of standard mixed model test statistics is small. However, our simulations suggest that for larger sample sizes, the effects of relatedness will become more pronounced, so we expect the two variance component model to become increasingly relevant as sample sizes increase.

While we are not aware of prior work in human genetics using two variance components to model effects of typed SNPs as well as additional phenotypic covariance from close relatives, other methods for combining these two sources of information for prediction have been proposed; however, these methods either use only a limited number of genome-wide significant SNPs [24] or use only limited information about family history [25]. Separately, several studies have applied different multiple variance component models to improve mixed model prediction and association in other ways. Widmer et al. [26] recently proposed a two variance component model that uses the standard GRM along with a GRM created from selected SNPs (as in FaST-LMM-Select [34]) that improves association power and calibration in family studies. (We note that while Widmer et al. observe that standard mixed model association is properly calibrated in their simulated family data sets, their simulations do not include untyped causal SNPs.) In another direction, Speed et al. [7] recently proposed a multiple variance component model that partitions SNPs into contiguous blocks, each used in a distinct variance component, and showed that this approach improves prediction accuracy. Incorporating a variance component modeling relatedness—either from pedigree, thresholding the GRM, or other approaches [35]—into these methods or recently proposed non-infinitesimal models for genetic prediction (e.g., weighted G-BLUP [6], BayesR [3, 10] and BSLMM [5]) is a possible direction for future research.

A challenge facing all genetic prediction methods is the very large sample sizes that will be required to achieve clinically relevant prediction accuracy [25, 36]. Indeed, in absolute terms, the prediction accuracy we achieved on real data sets of up to 8,000 samples was low, similar to other methods when applied to traits without large-effect loci [5, 6, 10]. Our simulations show that the two variance component approach we have proposed will maintain its relative improvement over standard BLUP as sample sizes increase; however, both of these methods face computational barriers at large *N*. A straightforward implementation of our two variance component method for prediction requires *O*(*N*^2^) memory and *O*(*N*^3^) time per REML iteration when estimating variance parameters as well as when computing predictions. These limitations could be overcome using a combination of rapid relationship inference [37], fast multiple variance component analysis (e.g., as implemented in BOLT-REML [38]), and iterative solution of the mixed model equations [39,40]. Similarly, the computational challenge of large-scale two variance component association analysis could potentially be addressed by extending fast iterative methods for mixed model association [16]. An alternative, computationally simple solution to inflation of association test statistics is LD Score regression [41]; however, this approach may incur slight deflation due to attenuation bias [16, 41].

We also note three additional limitations of our two variance component approach. First, the method is only applicable to data sets with related individuals for which genotypes are available for analysis; however, large human data sets of this type are now being generated (e.g., deCODE [42], 23andMe [43], and UK Biobank; see Web Resources). Second, the improved predictive performance of the two variance component approach is a function of the relatedness structure. Our parallel work in cattle has reported improved prediction accuracy using a two variance component model incorporating exact pedigree information [44] or breed information [45]; however, the two variance component model did not produce an improvement in analyses of Holstein dairy cattle (Table S8), perhaps due to the very small effective population size of this breed [46]. Third, our approach does not address case-control ascertainment. While many large family data sets are not ascertained for phenotype, investigating whether techniques employed by methods that do model ascertainment [8] can be integrated into our two variance component approach is a possible avenue for future work.

## Materials and Methods

### Standard mixed model for prediction

We begin by establishing notation and reviewing standard formulas for mixed model prediction (i.e., standard BLUP) and association testing using one variance component [1, 11, 27]. Let *N* be the number of individuals in the study and *M* be the number of genotyped SNPs. Denote phenotypes by *y*, fixed effect covariates by *X*, and normalized genotypes by *W*, all of which are mean-centered. We normalize each genotype by dividing by 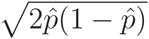, where 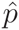 is the empirical minor allele frequency [18]. We model phenotypes using the following mixed model:

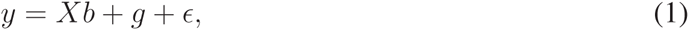

where *g* ∼ *N*(0, ∑_*g*_) is a random effect term modeling genetic effects, 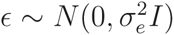 is a random effect modeling noise, and *b* is a vector of coefficients for the fixed effects. In the standard marker-based mixed model, we assume *g* = *W α* is a linear combination of genotyped SNPs, where *α* is an *M*-vector of iid normal SNP effect sizes (the infinitesimal model), so that

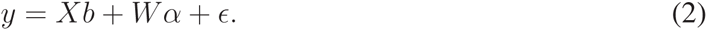

Then the genetic covariance satisfies 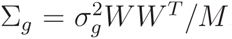, where *WW ^T^ /M* is the genetic relationship matrix and 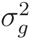 and 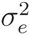 are variance parameters typically estimated using restricted maximum likelihood (REML) [47]. In pedigree-based models that do not use marker information, 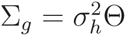, where Θ is the pedigree relationship matrix; again, 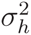 and 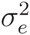 are estimated using REML.

These models naturally yield formulas for standard BLUP prediction [1]. Explicitly, if we denote training individuals (i.e., those with observed phenotypes) using subscript −*i* and denote test individuals (i.e., those with phenotypes to be predicted) using subscript *i*, predictions are given by

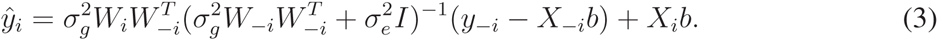

### Standard mixed model association test

To test a candidate SNP *w* for association with the phenotype *y*, we augment the marker-based model by including *w* as an additional fixed effect covariate:

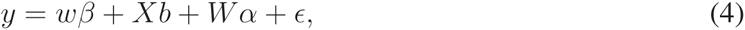

where *β* is the coefficient for the SNP *w* and we wish to test whether *β* ≠ 0. To do so, we estimate the variance parameters 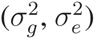 using REML and estimate the fixed effect coefficients (*β, b*) using maximum likelihood [27]. We may then compute the Wald statistic for testing *β* ≠ 0 as follows.

Let

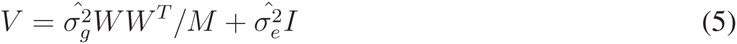

denote the total phenotypic covariance and let *Q* = [*w*; *X*] denote the combined fixed effects. Then 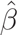 is equal to the first entry of (*Q^T^ V* ^−1^*Q*)^−1^*Q^T^ V* ^−1^*y* and var 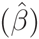 is equal to the first entry of (*Q^T^ V* ^−1^*Q*)^−1^. The Wald test statistic is given by

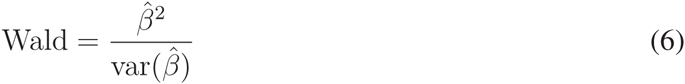

and is asymptotically *χ*^2^ distributed with 1 degree of freedom under the null distribution.

We make one slight modification to the above association test to avoid proximal contamination (i.e., masking of the association signal by SNPs included in the random effects term that are in linkage disequilibrium with the SNP being tested). Specifically, we use a leave-one-chromosome-out procedure in which when testing SNP *w*, we exclude all SNPs on the same chromosome as *w* from the genotype matrix *W* used to model random genetic effects [15, 33, 34]. Additionally, to save computation time, we fit variance parameters only once per left-out chromosome, reusing variance parameters from the null model [13] when computing approximate test statistics at all SNPs on the left-out chromosome [15, 16].

### Two variance component mixed model

Our use of a two variance component mixed model is motivated by the idea that in a sample containing related individuals, the pedigree relationship matrix (or an approximation thereof) can model additional heritable variance explained by untyped SNPs [17]. More precisely, consider expanding the marker-based model (2) to

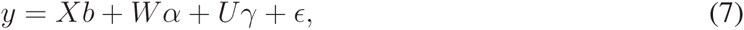

where *U γ* is the analog of *W α* for untyped SNPs *U*, so that the total genetic effect *g* = *W α* + *U γ*. Ideally, we would use this model and its augmentation for prediction and association testing, but *U* is unobserved. Because the BLUP and Wald statistic formulas only require *U U* ^*T*^, however, we can still improve upon the standard model (2) by using an approximation of *U U ^T^*. Letting *M_h_* denote the number of untyped SNPs, the matrix *U U ^T^/M_h_* is the realized relationship matrix from untyped SNPs. Assuming a fixed pedigree relationship matrix Θ, we have

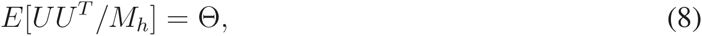

where the expectation is over possible realizations of genotypes passed down by descent (e.g., siblings share half their of genomes on average). When the study samples include close relatives, off-diagonal entries of Θ can be large, in which case these entries are good approximations of the corresponding entries of *U U ^T^ /M_h_* and hold additional information not fully harnessed by models that use only the usual GRM *W W ^T^ /M* from typed SNPs. Substituting Θ for *U U ^T^ /M_h_* gives the model

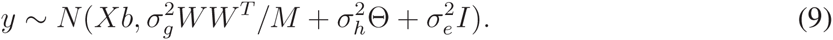

In our case, the pedigree relationship matrix Θ is also unavailable, so we need to make a further approximation in which we replace Θ with the estimator

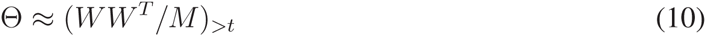

obtained from the usual GRM by keeping only those entries larger than a threshold *t* and setting all other entries to zero [17]. This approximation gives the model

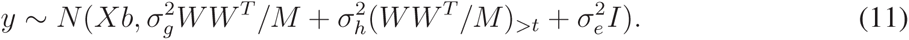

In theory, the optimal threshold *t* depends on *M, N*, and the amount of relatedness in the data set, but in our genetic prediction analyses using human data sets, we found that the results were robust to the choice of *t*, so we set *t* = 0.05. For association testing, we found *t* = 0.05 to generally be robust (and we expect this choice to be appropriate in human genetics settings), but in more extreme simulation scenarios in which we built the GRM from only a few chromosomes, we observed that higher thresholds were required to model relatedness accurately enough to produce well-calibrated statistics. We therefore optimize *t* in all association analyses (all of which we conduct using a leave-one-chromosome-out procedure [15, 33, 34]) using the following approach. For each chromosome *c* in turn, we choose *t* to minimize the deviation between the thresholded GRM 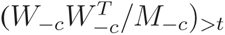 computed using all chromosomes but *c* and the GRM 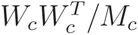 computed on-c the left-out chromosome *c*. We measure this deviation with the Frobenius norm

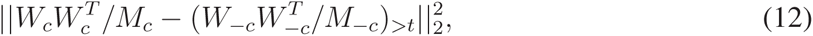

i.e., the sum of squared differences between matrix entries. Prediction and association testing proceed as before once the threshold *t* has been set: we estimate 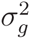, 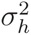, and 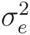 by REML to enable calculation of BLUP predictions, and for association testing, we again introduce an additional fixed effect term *wβ* for the SNP being tested and construct a Wald statistic. (Again, for computational efficiency, we apply a leave-one-chromosome-out procedure within which we reuse variance parameters fitted once per left-out chromosome.) We note that the computation of predictions 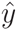 can no longer be expressed as a simple matrix-vector product between genotypes of testing individuals and a vector 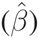 of SNP weights, as is the case for standard (one variance component) genomic BLUP. Instead, the formula for 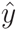 (given in Text S1) involves two terms, only one of which has the above form. We have not investigated the performance of prediction using the first term alone, as we expect that such a procedure, though computationally efficient, would yield suboptimal results.

### CARe and FHS data sets

We analyzed 8,367 African-American CARe samples from the ARIC, CARDIA, CFS, JHS and MESA cohorts with high-quality genotypes at 770,390 SNPs from an Affymetrix 6.0 array; the CARe data set and QC procedures used to obtain the sample and SNP sets we analyzed are described in refs. [21, 48]. We analyzed all samples in analyses of simulated phenotypes (using real genotypes); in analyses of real CARe phenotypes—BMI, height, HDL, and LDL, each available for 5,000–8,000 samples—we removed outlier individuals with phenotype values in the top or bottom 0.1%, individuals with age *<* 18, and individuals with missing age or sex; we then applied a Box-Cox transformation to remove skewness. We analyzed 7,476 FHS SHARe samples with high-quality genotypes at 413,943 SNPs from an Affymetrix 500K array and with BMI and height phenotypes available; the FHS data set and QC procedures are described in refs. [22, 23, 49].

### Genetic prediction: simulations with real genotypes

To assess the accuracy of genetic prediction methods, we simulated phenotypes based on genotypes from the CARe and FHS data sets, both of which are family studies containing many close relatives. Because the CARe individuals are admixed, we projected out the first 5 principal components (equivalent to including them as fixed effect covariates [47]) from genotypes and phenotypes in all analyses of both CARe and FHS data to avoid confounding from population structure [50]. We simulated phenotypes by generating causal effects for two subsets of SNPs: a set of *M* “observed SNPs,” which we used for both phenotype simulation and BLUP prediction, and a set of *M_h_* “untyped SNPs,” which we used for phenotype simulation but did not provide to prediction methods. In this simulation framework, the standard GRM built by MLM methods accurately models variation due to observed SNPs, but direct or inferred pedigree information is necessary to capture variation due to untyped SNPs. We generated effect sizes for observed and untyped

SNPs from independent normal distributions 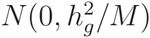 and 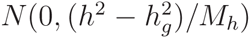, respectively, where 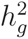 denotes heritability explained by observed SNPs and *h*^2^ denotes total narrow-sense heritability. To build phenotypes, we multiplied the simulated effect sizes with the genotypes and added random noise ∼ *N*(0, (1 − *h*^2^)). We used SNPs on chromosome 1 as untyped SNPs and used SNPs on varying subsets of chromosomes 2–22 as observed SNPs so as to simulate different values of *N/M* (which is a key quantity affecting performance of mixed model prediction [51] and association [15]) and thereby estimate projected performance at larger *N*. We used 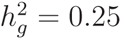 and *h*^2^ = 0.5 as typical values of these parameters [52].

We note that under the above setup, untyped SNPs are completely untagged by typed SNPs, whereas in real data, untyped SNPs may be partially tagged by typed SNPs. In either case, the phenotype can be written as a sum of “genetic value explained by typed SNPs,” “remaining genetic value,” and “environmental value” (with variance parameters 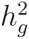, 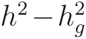, and 1 – *h*^2^ corresponding to the same covariance structures in either case), so we expect that our results are insensitive to this distinction (as evidenced by the fact that improvements in prediction accuracy in these simulations were corroborated by similar improvements in prediction accuracy on real phenotypes).

### Genetic prediction: simulations with simulated genotypes

To assess the potential performance of genetic prediction methods at extremely large sample sizes, we also simulated genotypes for sets of sib-pairs (relatedness = 0.5) with *M*=100 SNPs and *N/M*=10,20,…,100. We generated unlinked markers for simplicity by randomly generating minor allele frequencies uniformly in [0.05, 0.5] and sampling genotypes of unrelated individuals from a binomial distribution with the generated MAF. For sib-pairs, with probability 0.5, the pair shared an allele drawn randomly; otherwise, the alleles for the pair were drawn independently. (We ran this procedure twice per SNP to create diploid genotypes.) We simulated phenotypes as above.

### Genetic prediction: assessing performance on real phenotypes

To compare the predictive performance of the two variance component model versus standard BLUP on real phenotypes, we performed cross-validation studies in which we repeatedly selected 10% of the phenotyped samples (either CARe or FHS) as test data and used the remaining 90% of samples to train each predictor. For each train/test split *s*, we thus obtained a pair of observed prediction *r*^2^ values, 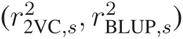. We then computed the relative improvement of the two variance component model over BLUP as

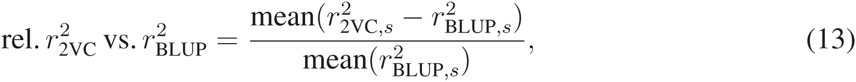

where the means are taken over the random splits *s*. We estimated the standard error of this quantity with the following expression:

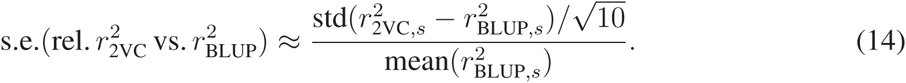

The numerator is the standard deviation of the per-split differences in *r*^2^ (across random 10% test sets *s*), which measures the variability in observed performance differences between the two methods when assessed on 10% of the data. We then divide by 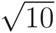 to account for the 10x larger sample size of the full data set, and finally, we normalize by the performance of BLUP to convert to the relative performance scale. This estimate is approximate due to the complexities of estimating variance under cross-validation (specifically, the slight overlap among different 10% test sets); in general, unbiased estimators of variance under cross-validation do not exist [53].

### Association testing: simulations with simulated genotypes

We conducted a suite of mixed model association simulations using genotypes simulated in a similar manner as above. We systematically varied the number of related individuals, the degree of relatedness, the number of markers *M* in the genome, and the SNP heritability *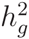* and total heritability *h*^2^ of the simulated trait. Specifically, we simulated sets of *N* = 1000 diploid individuals in which *N*_rel_ = 50, 125, 250, or 500 pairs of individuals were related and the rest were unrelated (leaving 900, 750, 500, or 0 unrelated individuals, respectively). Each pair of individuals shared a proportion *p* = 0, 0.1, 0.2, 0.3, 0.4, or 0.5 of their genomes in expectation. Additionally, we varied the number of markers *M* = 1,000, 5,000, 10,000, or 20,000. We generated unlinked markers as above; for pairs of related individuals, with probability equal to the relatedness *p*, the pair shared an allele drawn randomly; otherwise, the alleles for the pair were drawn independently. (As above, we ran this procedure twice per SNP to create diploid genotypes.) We further generated 100 additional candidate causal SNPs and 500 candidate null SNPs (at which to compute association test statistics) in the same way. We used an infinitesimal model to generate the phenotype: that is, we generated effect sizes for the observed SNPs from *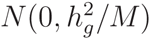*. We also generated effect sizes for the candidate causal SNPs from 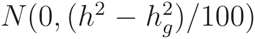. Because these SNPs are distinct from the *M* SNPs used for model-building, they effectively served as untyped causal loci. Finally, we formed the phenotype by multiplying the effect sizes with the genotypes and adding independent noise distributed as *N*(0, (1 – *h*^2^)*I*).

### Association testing: simulations with real genotypes

We also assessed mixed model association methods in simulation studies using simulated phenotypes based on genotypes from the CARe and FHS data sets. To avoid proximal contamination [15, 33, 34], we tested SNPs on chromosomes 1–2 for association and used *M* “observed SNPs” on subsets of chromosomes 3–22 to compute GRMs, varying the number of chromosomes used to vary *N/M*. We generated quantitative phenotypes in which observed SNPs collectively explained 25% of variance and 250 causal SNPs from chromosome 1 explained another 25% of variance; all SNPs on chromosome 2 were null SNPs.

## Web Resources

UK Biobank, http://www.ukbiobank.ac.uk/

## Acknowledgments

We are grateful to N. Zaitlen, B. Vilhjalmsson, S. Rosset, and H. Johnsen for helpful discussions. This research was supported by NIH grants R01 HG006399 and R01 GM105857 and NIH fellowship F32 HG007805.

